# Transcriptomic Mutational Profiling of Gastric Adenocarcinoma in Northern Brazil

**DOI:** 10.1101/2025.09.15.676424

**Authors:** Rubem Ferreira da Silva, Diego Pereira, Mateus Silva Tavares, Daniel de Souza Avelar, Jéssica Manoelli Costa da Silva, Ronald Matheus da Silva Mourão, Valéria Cristiane Santos da Silva, Juliana Barreto Albuquerque Pinto, Samir Mansour Casseb, Samia Demachki, Tatiane Neotti, Ana Karyssa Mendes Anaissi, Williams Fernandes Barra, Amanda Ferreira Vidal, Geraldo Ishak, Paulo Pimentel de Assumpção, Rommel Mario Rodriguez Burbano, Fabiano Cordeiro Moreira

## Abstract

Gastric cancer (GC) remains among the neoplasms with the worst prognosis, partly due to its biological heterogeneity and the scarcity of robust biomarkers. The characterization of mutational profiles from transcripts can reveal specific tumor signatures and point to therapeutic targets. This study investigated the landscape of mutations expressed in 102 samples of gastric adenocarcinoma from northern Brazil, which were sequenced using NGS. Readings were aligned to the reference genome using STAR (two-pass mode), and variants were called with GATK and VarDict. Annotations and impact predictions were generated using VEP, SIFT, and PolyPhen. We identified >90,000 variants; among the most frequently mutated genes, FTH1 stood out. The mutational profile was described using maftools, and signatures were inferred with MutationalPatterns. We observed a predominant distribution of SNVs, with C>T transitions as the most common event, in addition to patterns compatible with signatures related to replication damage and DNA repair. To mitigate biases inherent to RNA-seq, we applied filters for coverage, strand bias, and RNA editing hotspots. Together, the data outline a regional landscape of mutations expressed in GC and reinforce the usefulness of the transcriptome for prioritizing biomarkers and functional hypotheses that may guide genomic validations and subsequent clinical studies.

## INTRODUCTION

Mutational processes are a cumulative phenomenon in somatic cells, with most mutations being silent and having no functional impact (FORMA, 2020). However, some mutations can be triggered by factors such as UV radiation, chemicals, or endogenous errors in DNA repair, affecting vital cellular functions (MARTICORENA and CAMPBELL, 2015). Understanding these mutations is crucial, especially in the context of gastric carcinogenesis, where the accumulation of mutations in driver genes affects cellular processes such as DNA repair and signal transduction (FORMA, 2020).

Despite remarkable advances in cancer treatment, the identification of reliable biomarkers and the ineffectiveness of targeted therapies in most cases remain persistent challenges in clinical practice (PATEL, 2017). Although next-generation sequencing (NGS) technologies and robust bioinformatics tools have revolutionized genomic analysis, allowing for more detailed and thorough investigation of molecular data (HOLTSTRATER, 2020), progress in the development of personalized treatments still faces significant barriers. One of the most critical issues is the underrepresentation of ethnically diverse populations in genomic studies (SIRUGO, 2019).

In this scenario, RNA-seq emerges as a promising tool, as it bridges the gap between DNA and protein, offering greater clarity and predictability for precision oncology. The variants detected by RNA-seq are directly linked to the transcriptome, allowing immediate understanding of the functional impacts of mutations and revealing whether they are associated with positively or negatively regulated genes or pathways (LI et al., 2025; Bollas et al., 2025).

## METHODOLOGY

### Sample Characterization and Ethical Considerations

As part of the main project to which this proposal belongs, 102 tumor tissue samples were collected from patients diagnosed with gastric adenocarcinoma. The samples were obtained from the João de Barros Barreto University Hospital (HUJBB) and the Ophir Loyola Hospital.

All participants were fully informed about the research objectives, and samples were collected only after obtaining informed consent by signing the Free and Informed Consent Form (FICF). The use of all samples and the execution of this study were approved by the Research Ethics Committee of the João de Barros Barreto University Hospital, under protocol number CAAE 47580121.9.0000.5634. For the collection of tumor tissue, 0.5 cm fragments were surgically resected. These fragments were collected immediately after gastric resection, preserved in RNAlater for transport, and stored in a freezer at -80 °C.

### Ethics statement

This study was approved by the Ethics and Research Committee of João de Barros Barreto University Hospital (approval number: 47580121.9.0000.5634) and was conducted in accordance with the principles outlined in the Declaration of Helsinki. Participant recruitment and sample collection were carried out between July 2, 2022, and July 6, 2023. Before enrollment, all participants received detailed information about the study’s objectives, potential benefits, risks, and possible harms, ensuring a thorough understanding of the research. Written informed consent was voluntarily obtained from all participants prior to their inclusion in the study.

### Clinical Characterization of Patients

The medical records and other clinical data of the patients investigated were reviewed for the presence of gastric adenocarcinoma, as well as for gender, age, Lauren histological subtype, TNM pathological staging, administration of perioperative chemotherapy regimens with 5-Fluorouracil, Leucovorin, Oxaliplatin, and Taxane (FLOT), and mortality. TNM staging was determined based on the 8th edition of the AJCC TNM classification for gastric cancer.

### Total RNA extraction

Initially, approximately 30 mg of tissue from each sample was macerated. Next, 1 mL of TRIZOL® reagent was added to the processed tissue for extraction. TRIZOL® reagent (Thermo Fisher Scientific) was used to preserve the integrity of cellular RNA and promote cell lysis. After centrifugation at 13,000 rpm for 10 minutes at 4 °C, RNA was recovered by precipitation with isopropyl alcohol. The precipitated RNA (total RNA) was washed with ethanol (EtOH), air-dried at room temperature, and subsequently analyzed for integrity and concentration using the Qubit 2.0 Fluorometer (Thermo Fisher Scientific), NanoDrop ND-1000 (Thermo Fisher Scientific), and the 2200 TapeStation System (Agilent). The ideal criteria for total RNA integrity included values between 1.8 and 2.2 for the A260/A280 ratio, greater than 1.8 for the A260/A230 ratio, and an RNA Integrity Number (RIN) of 5 or higher.

### cDNA Library Construction

For library construction, the TruSeq Stranded Total RNA Library Prep Kit with Ribo-Zero Gold (Illumina, USA) was used according to the manufacturer’s instructions. In preparing the library, 1 μg of total RNA was used per sample, in a final volume of 10 μL. After constructing the libraries, a new integrity assessment was performed using the 2200 TapeStation System. At the end of the process, a band of ∼260 base pairs was observed.

### NGS Sequencing and Read Quality Control

The previously constructed cDNA libraries were loaded onto the Illumina NextSeq sequencing system and sequenced using the paired-end method (reads were generated from both the forward and reverse strands of the cDNA). The NextSeq 500 MID Output V2 kit (Illumina, 150 cycles) was used to process the libraries under the conditions specified by the manufacturer. Initially, read quality was assessed using FastQC. Next, adapters and low-quality reads were removed using Trimmomatic v0.39, with a quality value (QV) set above 25.

### Preprocessing and Variant Calling

The fastq files were aligned using the STAR v2.7.11b tool (Dobin, 2013) in two-pass mode using the default configuration. The BAM files were processed following the Genome Analysis Toolkit (GATK) protocols. Initially, they were sorted using SortSam, and duplicates were marked with MarkDuplicates. Next, the SplitNCigarReads tool was used to process reads spanning splicing between exons and introns. After base recalibration steps to ensure sequence quality, variant calling was performed with the VarDict tool, generating VCF files with the identified variants.

### Filtering and Impact Analysis

The generated VCF files underwent a rigorous filtering process. Variants with a depth of reading (DP) lower than 10 and those with allele frequencies between 45-55% and 95-100% were removed to mitigate the inclusion of heterozygous and germline homozygous variants (Jessen, 2021). Only variants that received a “PASS” signal from VarDict were retained. The variants were then annotated using the Ensembl Variant Predictor (VEP), and synonymous mutations were excluded. For functional impact prediction, the SIFT and PolyPhen-2 tools were used, classifying variants with SIFT scores between 0 and 0.05 as deleterious.

### Mutational Profile and Signature Analyses

To characterize the mutational profile, VCF files were converted to MAF format using VCF2MAF. Analyses were conducted in the R environment using the Maftools and MutationalPatterns packages (Manders et al., 2022). Maftools (Mayakonda et al., 2018) was used to visualize the mutational landscape through oncoplots, determine the mutation burden per sample, and identify the most frequently mutated genes. Mutational signatures were extracted with MutationalPatterns, using the COSMIC database (v3) to identify the most representative signatures.

## RESULTS

In this study, 102 gastric cancer tumor samples were analyzed, focusing on aspects such as mutations per sample, frequently altered genes, co-occurrence effects, and the most representative mutational signatures.

### Summary plot of key information from the dataset

Figure 1 presents an overview of the identified mutations, consolidating several analyses into six panels. The classification of variants (upper left panel) shows that ‘Missense_Mutation’ mutations are the most frequent, followed by reading frame deletions. In terms of variant type (top center panel), single nucleotide mutations (SNPs) are the most prevalent, surpassing deletions and insertions. Analysis of nucleotide substitutions (top right panel) reveals a high prevalence of C>T mutations, a pattern that may indicate specific mutational processes. The distribution of mutations per sample (lower left panel) shows a wide variation in mutational burden, with a median of 427.5 mutations per sample. Finally, the 10 most mutated genes (lower right panel) are highlighted, with the FTH1 gene showing the highest mutation frequency (47%), followed by MPDU1 (41%) and TXNIP (40%).

**Figure 1.**
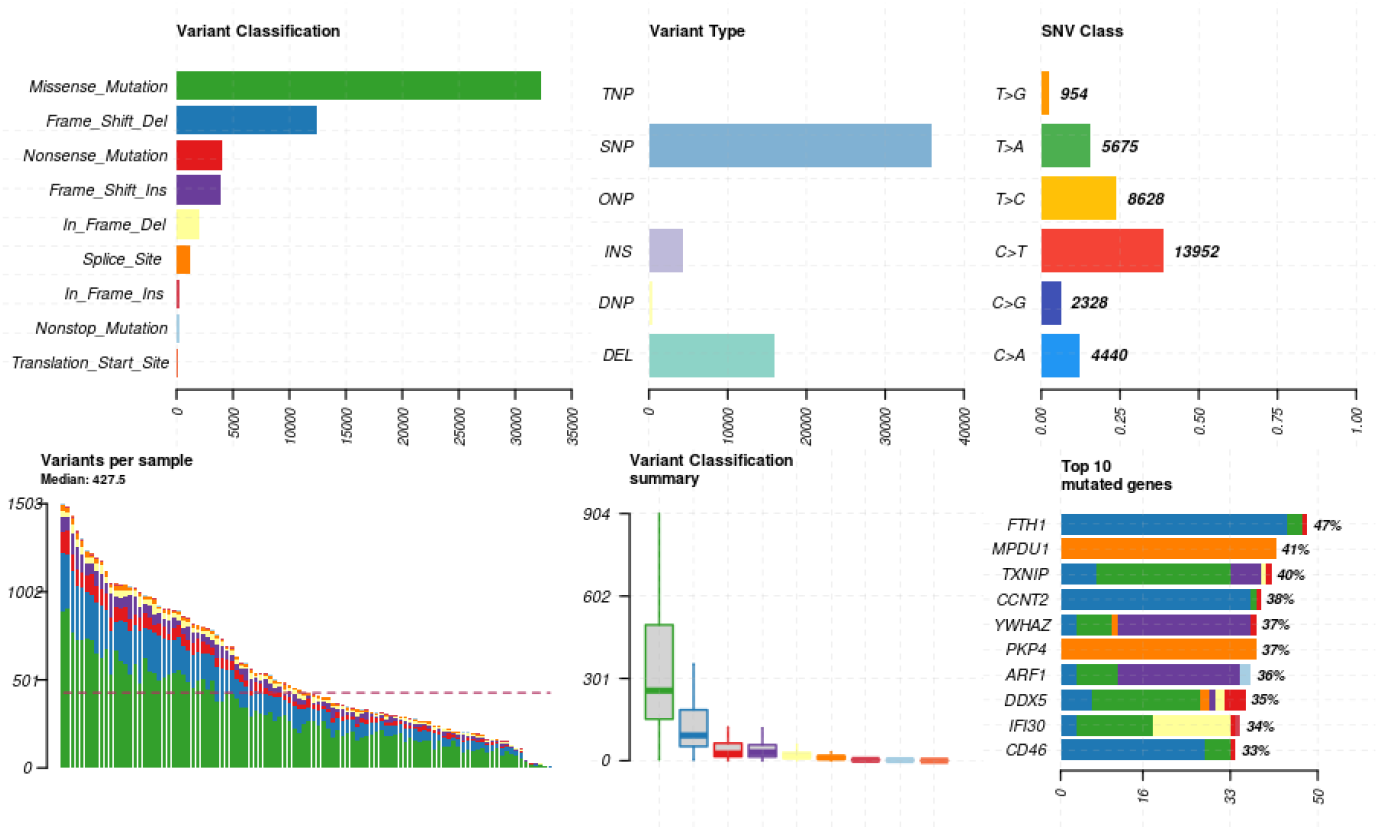
Visualizing mutation summary. This maftools plot shows a summary of the MAF file. Highlighting the most mutated genes, SNV class, and variant classification distributions within a tumor type.

### Mutations per absolute chromosome and normalized by chromosome size

Figure 2 details the distribution of mutations across chromosomes and by base substitution type. Panel A shows the absolute distribution of mutations, where larger chromosomes, such as chr1, exhibit the highest number of mutations. To correct for the influence of chromosome size, Panel B normalizes the data per million base pairs, revealing that chr19 has the highest mutational density. Panel C summarizes the total number of mutations, highlighting that C>T substitutions are the most prevalent across the genome, followed by G>A and T>C. Together, these panels show that mutation frequency is not uniform and that C>T mutations are frequent in the mutational profile.

**Figure 2.**
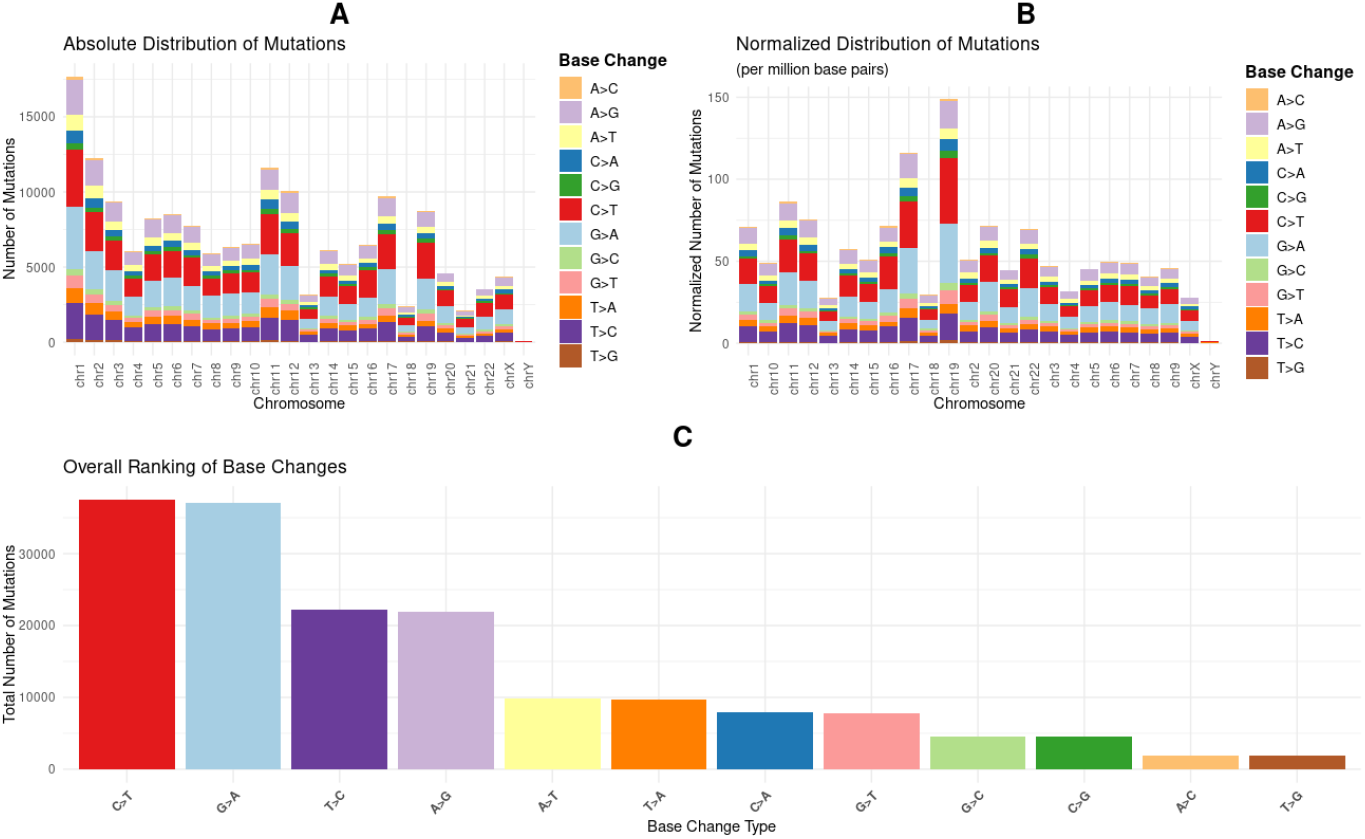
illustrates the distribution of mutations in human chromosomes. Panel A shows the absolute distribution of mutations by chromosome, while panel B shows the normalized distribution. Panel C ranks the total frequency of each mutation type.

**Figure 4.**
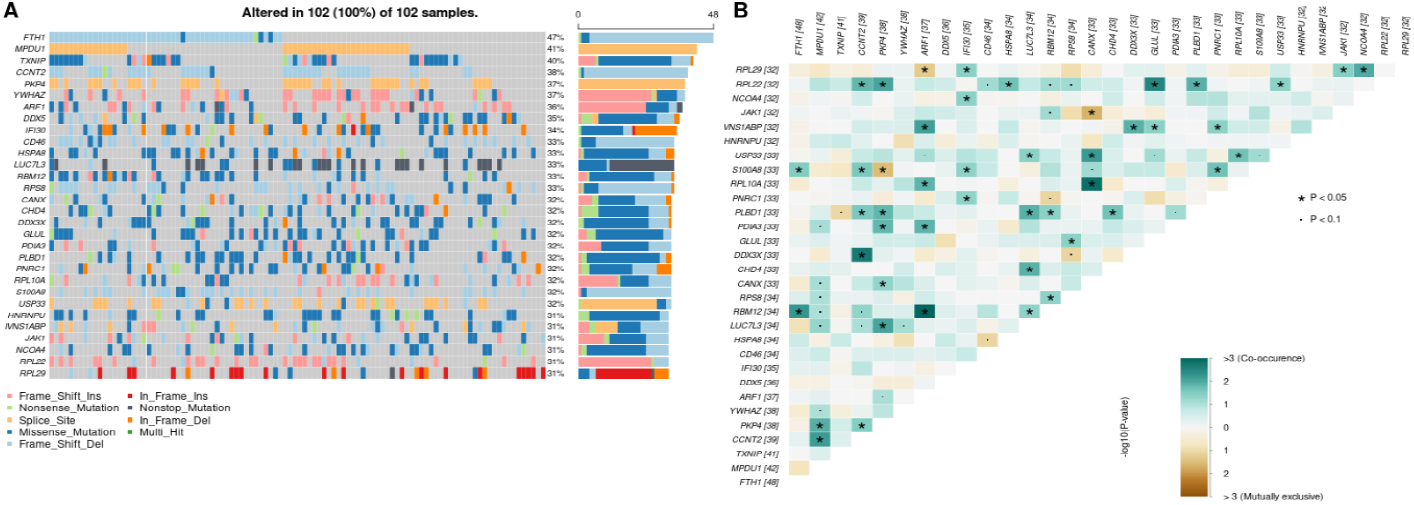
combines two graphs to show the relationship between genetic mutations. Panel (A) on the left is an oncoplot illustrating the presence of mutations in the 30 most frequently mutated genes in the 102 samples. The panel on the right (B) is a matrix showing the co-occurrence (green) or mutual exclusion (orange) of these mutations. Together, the graphs describe the mutation pattern and frequency of co-occurrence in a cohort of samples. The symbols indicate statistical significance for the Mann-Whitney U test: *P<0.05; **P<0.01.

### Oncoplot and co-occurrence effect

Figure 2 shows the mutation profile of the most altered genes, showing their frequency and co-occurrence. Panel A (oncoplot) illustrates the mutations in the 30 most frequent genes, highlighting the FTH1 gene as the most mutated (47%). Panel B (co-occurrence matrix) shows the associations between these genes. Although there are some statistically significant co-occurrences (marked with asterisks), most genes do not mutate together. This suggests that the mutational profile is heterogeneous among samples.

### Oncoplot of the most frequently mutated genes in gastric cancer

Figure 5 presents an oncoplot summarizing the mutation profile of the most biologically relevant genes, with the aim of highlighting alterations in genes already well established in the literature. Each row of the graph represents a gene, and each column corresponds to an individual sample. The colored boxes indicate the presence and type of mutation, as shown in the legend. The ARID1A, RHOA, and CTNNB1 genes are the most frequently mutated, with frequencies of 23%, 23%, and 21%, respectively.

**Figure 5.**
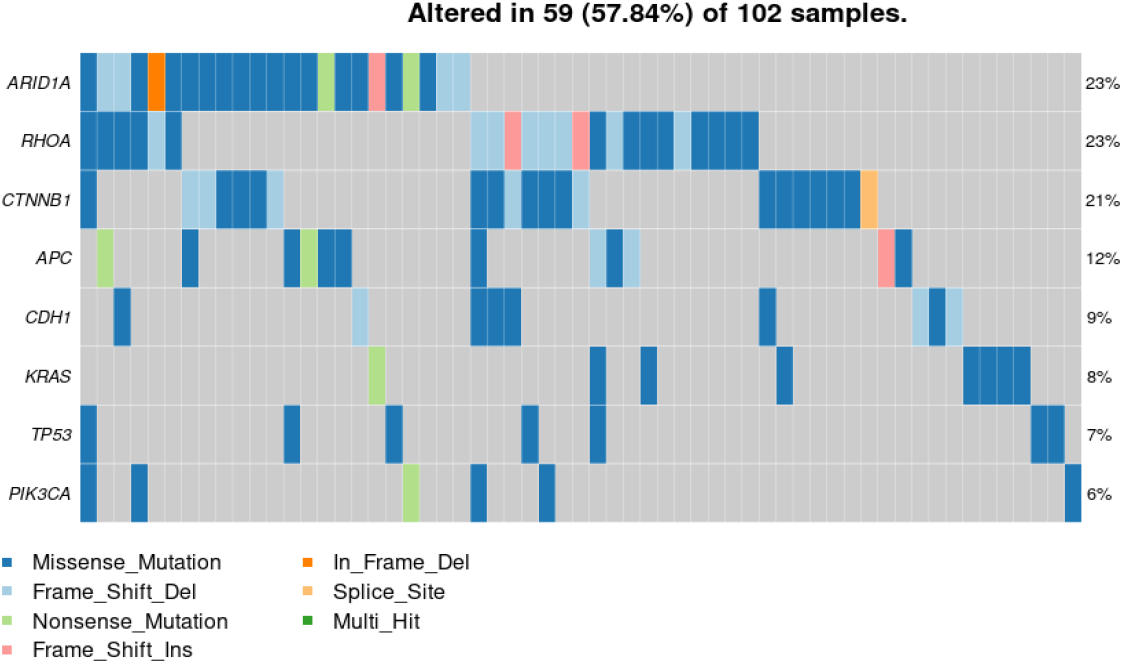
The oncoplot illustrates the frequency and type of mutations in the most commonly altered genes in 102 gastric cancer samples. As indicated, 59 (57.84%) of the samples had at least one alteration in one of the analyzed genes.

### Contribution of mutational signatures

Figure 6 details the contribution of mutational signatures to the observed mutations. Panel A illustrates the relative contribution of each signature per sample, showing that the heterogeneity between samples is significant. The overall analysis (Panels B and C) highlights that the SBS5 and SBS3 signatures are the most prevalent. The predominance of SBS5 suggests the action of a “clock” process, while SBS3 is associated with deficiencies in DNA damage repair. These results indicate that the mutational landscape is not random and can be attributed to specific biological processes.

**Figure 6.**
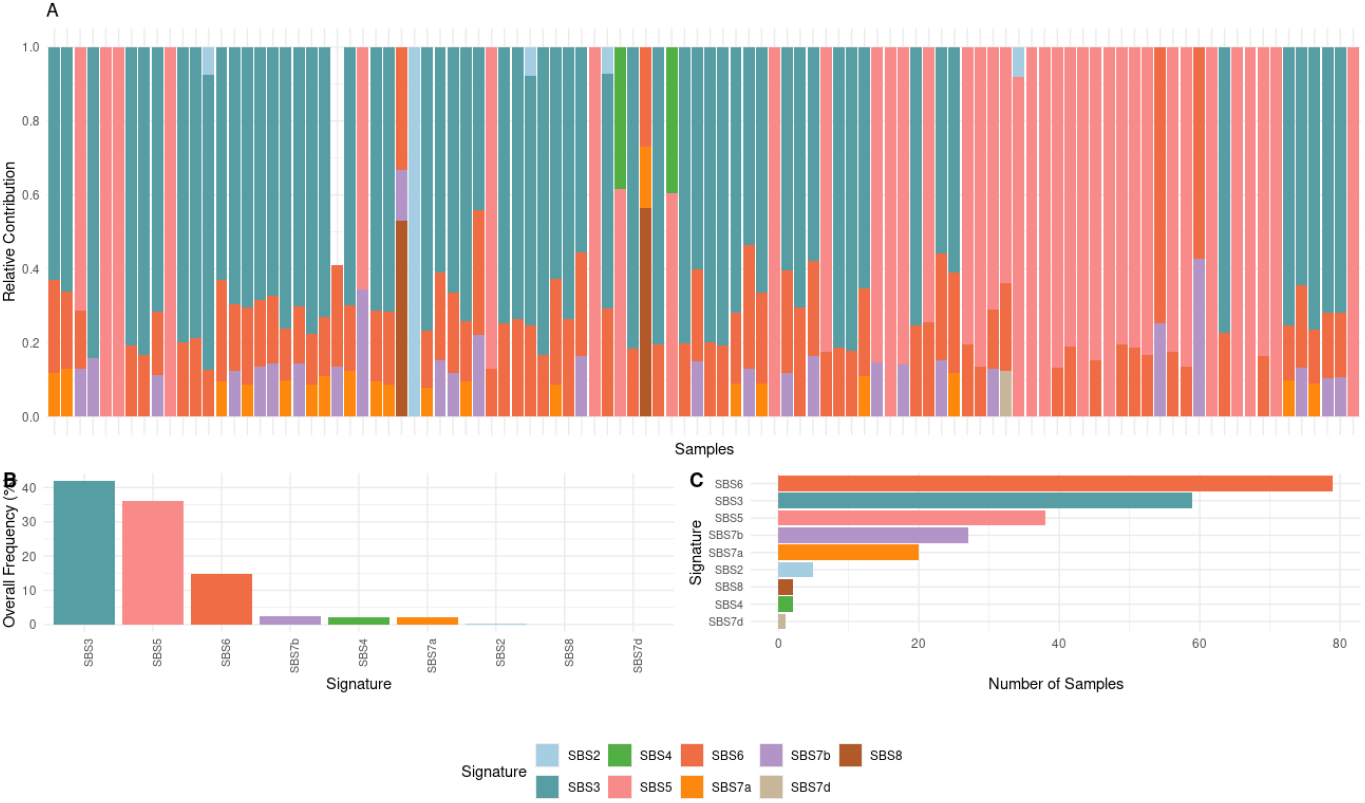
Mutational signatures, which reflect the biological processes that cause mutations, are detailed in the lower panel. Panel A shows the relative contribution of each mutational signature per individual sample. In Panel B, the overall frequency of each signature in the cohort is presented. Panel C indicates the total number of samples where each signature was detected. The color legend for the mutational signatures is provided at the bottom.

## DISCUSSION

Analysis of the mutational profile showed a predominance of missense mutations, followed by frameshift deletions and nonsense mutations, indicating a significant impact on protein structure. The most frequent variant type was SNP, with emphasis on C>T substitution, a pattern commonly associated with mutagenesis processes in gastric cancer. The mutational burden varied widely among samples, with a median of 427.5 variants per patient, reflecting relevant intratumoral heterogeneity. Among the most frequently mutated genes were FTH1 (47%), MPDU1 (41%), TXNIP (40%), and CCNT2 (38%).

The chromosomal distribution of mutations showed a higher absolute number on chromosomes 1, 2, and 17, partly due to their size and gene density. After normalization by length, chromosomes 17 and 19 exhibited disproportionate mutational burden, suggesting hotspots of instability. In the context of gastric cancer, chromosome 17 concentrates critical genes (TP53, ERBB2, BRCA1) associated with worse prognosis and targeted therapies (Cancer Genome Atlas Research Network, 2014; Cristescu et al., 2015). In the global ranking of substitutions, the predominance of C>T, followed by T>C and G>A, indicates endogenous processes (cytosine deamination and MMR failures), consistent with COSMIC signatures (SBS3, SBS5, SBS6) often implicated in carcinogenesis.A, indicates endogenous processes (cytosine deamination and MMR failures), consistent with COSMIC signatures (SBS3, SBS5, SBS6) frequently implicated in gastric carcinogenesis (Alexandrov et al., 2020; Alexandrov et al., 2013; Zou et al., 2021).

In the oncoplot, FTH1, TXNIP, DDX5, CHD4, and JAK1 are the most frequently mutated genes, connecting iron homeostasis/ferroptosis (FTH1) (Liu et al., 2022), redox imbalance with prognostic potential (TXNIP) (Lim et al., 2012; Deng et al., 2023), post-transcriptional control/chromatin remodeling (DDX5) (Wang et al., 2023), epigenetic regulation linked to EMT/WNT (CHD4) (Shi et al., 2025), and JAK/STAT hyperactivation (JAK1) mediated by circFCHO2 and HOXA10 (Khanna et al., 2015; Zhang et al., 2022; Chen et al., 2019). The significant co-occurrences of RBM12–ARF1 and RPL10A–CANX suggest a coupling between transcript regulation and vesicular trafficking (de Las Heras-Rubio et al., 2021; Wang et al., 2024) and dependence on translational capacity/glycoprotein folding in the ER (Oakes, 2016), pointing to tumor adaptation sustained by UPR/proteostatic stress (Wang et al., 2019).

In 102 samples, 57.8% had mutations in the genes evaluated. ARID1A and RHOA were the most frequent (∼23% each). ARID1A, from the SWI/SNF complex, is associated with genomic instability, EBV positivity, and higher PD-L1, suggesting a potential benefit from immunotherapy (Li et al., 2019; Angelico et al., 2024). RHOA, typical of the diffuse subtype, is related to cytoskeletal dynamics and loss of tissue cohesion (Zhou et al., 2014; Zhang et al., 2019; Nam et al., 2019). Alterations in CTNNB1 (∼21%) and APC (∼12%) indicate Wnt/β-catenin activation (Han et al., 2024; McGowan et al., 2023; Yan et al., 2022). Mutations in CDH1 (∼9%) reinforce the loss of epithelial adhesion in diffuse (Wang et al., 2024; Lim et al., 2023). KRAS (∼8%) and PIK3CA (∼6%) suggest MAPK and PI3K/AKT activation, with therapeutic implications (Yuan et al., 2024; Morgos et al., 2024). The frequency of TP53 (∼7%) was lower than in large cohorts, possibly due to subtype composition and/or methodological aspects, reflecting tumor heterogeneity (Cristescu et al., 2015; Oliveira et al., 2024).

Regarding the signatures found, SBS3 reflects homologous recombination deficiency (HRD), often linked to BRCA1/2 and HR genes, recognized in solid tumors, including gastric tumors (Alexandrov et al., 2020; Zhao et al., 2021). SBS6 translates microsatellite instability (MSI) due to MMR deficiency, a marker with predictive value for immunotherapy (Alexandrov et al., 2015; Liu et al., 2018). SBS5, ubiquitous and with low specific etiological impact, reflects aging/chronic endogenous processes (Alexandrov et al., 2020). Thus, the predominance of SBS3 and SBS6 suggests that HRD and MMR are central determinants of this mutational landscape, while minority signatures (SBS7, SBS8, and SBS2) possibly reflect more restricted or contextual processes.

## CONCLUSION

The identification of variants expressed by RNA-seq reveals particularities of the mutational landscape in patients from the North region with gastric adenocarcinoma, by integrating mutation types, variant frequency per gene, and mutational profiles/signatures.

## Acknowledgment

The authors would like to thank the Oncology Research Center, the Human and Medical Genetics Laboratory, and the Anatomical Pathology Laboratory at João de Barros Barreto University Hospital (HUJBB – UFPA) for their invaluable technical and laboratory support. Our gratitude also goes to the High-Performance Computing Center (CCAD) at the Federal University of Pará for access to the Apollo 2000 cluster, which was crucial for our analyses.

